# Significant Variability exists in the Toxicity of Global Methicillin-resistant *Staphylococcus aureus* Lineages

**DOI:** 10.1101/2021.07.16.452633

**Authors:** Maisem Laabei, Sharon J. Peacock, Beth Blane, Sarah L. Baines, Benjamin P. Howden, Timothy P. Stinear, Ruth C. Massey

## Abstract

*Staphylococcus aureus* is a major human pathogen where the emergence of antibiotic resistant lineages, such as methicillin-resistant *S. aureus* (MRSA), is a major health concern. While some MRSA lineages are restricted to the healthcare setting, the epidemiology of MRSA is changing globally, with the rise of specific lineages causing disease in healthy people in the community. In the past two decades, community-associated MRSA (CA-MRSA) has emerged as a clinically important and virulent pathogen associated with serious skin and soft-tissue infections (SSTI). These infections are primarily toxin driven, leading to the suggestion that hyper-virulent lineages/multi-locus sequence types (STs) exist. To examine this, we compared the toxic activity of 475 MRSA isolates representing five major MRSA STs (ST22, ST93, ST8, ST239 and ST36) by employing a monocyte-macrophage THP-1 cell line as a surrogate for measuring gross cytotoxicity. We demonstrate that while certain MRSA STs contain highly toxic isolates, there is such variability within lineages to suggest that this aspect of virulence should not be inferred from the genotype of any given isolate. Furthermore, by interrogating the accessory gene regulator (Agr) sequences in this collection we identified several Agr mutations that were associated with reduced toxicity. Interestingly, the majority of isolates that were attenuated in toxin production contained no mutations in the *agr* locus, indicating a role of other undefined genes in *S. aureus* toxin regulation.

## Introduction

*Staphylococcus aureus* is responsible for a wide array of diseases ranging from superficial skin infections to severe, life-threatening cases of pneumonia and bacteraemia (1). The emergence of antibiotic resistant lineages including methicillin-resistant *S. aureus* (MRSA) has complicated treatment. For decades the circulating MRSA lineages appeared to be limited to causing infections within healthcare settings in patients with predisposing conditions who were susceptible to infection (2). However, in the 1990s distinct MRSA lineages, unrelated to earlier circulating MRSA lineages, started to emerge outside of healthcare settings to cause infections in otherwise healthy individuals (2, 3). Community-associated MRSA (CA-MRSA) isolates result in similar clinical manifestations, such as severe skin and soft-tissue infections (SSTIs), despite broad genetic diversity among CA-MRSA lineages (4). Understanding the differences between the healthcare restricted MRSA lineages and the more recently emerged CA-MRSA lineages has been the focus of much attention (3).

Molecular and epidemiological studies of CA-MRSA isolates have identified multiple putative virulence factors, namely toxins, associated with the hyper-virulent phenotype characteristic of CA-MRSA isolates. The majority of CA-MRSA isolates harbour *lukSF-PV* which encodes the bi-component pore-forming leucocidin (4). Human neutrophils have been shown to be highly susceptible to PVL-mediated lysis due to the expression of complement receptors, C5aR and C5L2, which are required for PVL binding (5). Importantly, the role of PVL in pathogenesis is likely dependent on the site of infection; a clear role is observed in a rabbit necrotizing pneumonia model (6), but the role of this toxin in dermonecrosis is less evident (7, 8). Aside from PVL, over-expression of core genome virulence determinants, notably α-haemolysin (Hla) and α-type phenol-soluble modulins (α-PSMs), has been hypothesised to significantly contribute to the enhanced virulence of CA-MRSA isolates (9–12). Hla is the prototypical β-barrel pore-forming cytotoxin (13). Multiple studies utilising serological and animal models of infection have indicated a prominent role for Hla in the pathogenesis of disease (14). The α-PSMs are characterised as small amphipathic α-helical peptides that efficiently lyse numerous cell types, independent of cell specific receptor (11, 15). As with Hla, α-PSMs significantly contribute to virulence in a murine infection models of bacteraemia and skin lesions (11).

Toxin regulation in *S. aureus* is governed primarily by the accessory gene regulator (*agr*), which employs a cell density dependent, quorum sensing system to upregulate a suite of secreted virulence factors such as toxins and downregulate surface binding proteins (16). Accordingly, the *agr* system plays a central role in the development of a range of *S. aureus* infections, most notably in CA-MRSA skin infections (17). Furthermore, transcriptomic data indicate enhanced *agr* regulation of important toxins (PVL, Hla and α-PSM), which are frequently associated with CA-MRSA virulence (17). Together, these studies have contributed to the frequent use of the term ‘hyper-virulent’ when referring to CA-MRSA lineages.

Recently, studies of toxicity at a population level have reported that this phenotype varies widely within individual MRSA multi-locus sequence types (STs) (18), suggesting that the virulence of an individual MRSA isolate should not be inferred from its genotype (18). Non-cytolytic clinical isolates are commonly referred to as ‘Agr dysfunctional’, due to the frequency at which mutations occur within the sensor kinase (AgrC) or response regulator (AgrA) encoding genes, impacting significantly on toxin expression. To examine this in greater detail, and across a globally representative collection of isolates, we focussed on a collection of 475 MRSA isolates representing five major MRSA STs (ST22, ST93, ST8, ST239 and ST36; see Supp. Table 1 for further details) collected from Europe, North and South America, Asia and Australia, and examined their toxicity based on their ability to lyse human cells. As no single cell line exists that is susceptible to all of the toxins secreted by *S. aureus*, we use the THP-1 monocyte-macrophage cell line, which based on our empirical evidence, is susceptible to the widest range of *S. aureus* toxins, expressing receptors for PVL (5) and Hla (19) and susceptible to δ-haemolysin and α -PSMs (20). Our findings indicate that toxicity varies significantly within STs and therefore lineage should not be used as a metric to infer virulence.

In addition, we have identified novel Agr mutations associated with attenuated toxicity and confirm that multiple isolates with reduced toxicity have no mutations within the Agr operon, indicating the existence of other undefined toxin regulating genes in *S. aureus*.

## Materials and Methods

### Bacterial isolates and growth conditions

A list of the MRSA clinical isolates used in this study can be found in Supplementary Table 1. *S. aureus* isolates were grown overnight in 5 ml of Tryptic-Soy Broth (TSB; Sigma) in a 30 ml glass tube at 37°C with shaking at 180 rpm. Overnight cultures were used to inoculate 5 ml of fresh TSB at a dilution of 1:1000 and incubated for 18h at 37°C with shaking at 180 rpm. *S aureus* toxin containing supernatants were harvested from 18h cultures by centrifugation at 14,600 rpm for 10 min. All clinical isolates have been genome-sequenced as described previously (18, 20–24). Paired-end reads for these isolates were mapped to the following reference isolates: ST22 (HO 50960412 (24)), ST93 (JKD6159 (21, 22)), ST8 (USA300 strain LAC (20)), ST239 (TW20 (18, 23)) and ST36 (MRSA252 (24)). The accession numbers for the sequence data for each of these isolates are listed in the indicated reference.

### THP-1 cell culture

THP-1 monocyte-macrophage cell line (ATCC#TIB-202) was routinely grown in suspension in 30 ml of RPMI-1640 medium (Gibco: 11340892), supplemented with 10% heat-inactivated fetal bovine serum (FBS) (Sigma: F7524), 1 μM L-glutamine, 200 units/ml penicillin and 0.1 mg/ml streptomycin (Sigma: G6784) at 37°C in a humidified incubator with 5% CO_2_. THP-1 cells were routinely viewed microscopically and sub-cultured every 3-4 days. For use in cytotoxicity assays, cells were collected by centrifugation at 1000 rpm for 5 min at room temperature and resuspended to a final density of 1-1.2 × 10^6^ cells/ml in tissue grade phosphate buffered saline (Gibco). Cell viability was analysed using the Guava ViaCount reagent (Luminex) and easyCyte flow cytometry, typically yielding >95% viability following THP-1 collection.

### Cytotoxicity assay

The cytotoxicity assay was optimised previously (20). Briefly, to evaluate *S aureus* toxicity, 20 μl of harvested supernatant (either used as 100% or diluted to 70%, 30% or 10% in TSB) was incubated with 20 μl of washed THP-1 cells for 12 min at 37°C under static conditions. Cell death was quantified using the Guava ViaCount reagent and easyCyte flow cytometry according to manufacturer’s instructions. The toxicity of each isolate was measured with two technical repeats and three biological repeats with error bars indicating the standard deviation (SD). LAC (a highly toxic CA-MRSA isolate) supernatant and TS broth were used as positive and negative controls, respectively.

### Statistical analysis

A one-way ANOVA with Tukey’s multiple comparison test was used to examine differences between experimental results (GraphPad Prism v9.0), where a *p* value <0.05 was considered to be statistically significant.

## Results and Discussion

### S. aureus *toxicity is highly variable both between and within sequence types*

In this study we compared the toxicity profiles of previously published collections of MRSA isolates (i.e. sequence type (ST) 8 (20), ST22 and ST36 (24); n=330) with newly derived toxicity profiles (ST93 and ST239; n=145) to provide a comprehensive and globally distributed picture of toxin expression by *S. aureus*. What was immediately apparent was that the individual STs contained numerous isolates that were either extremely toxic (100% cell death) or non-toxic (0% cell death), and could not be reliably assayed under the same conditions. As a result, we used the supernatant of two of the STs (ST239 and ST36) neat (100%) and for the other three STs (ST22, ST93 and ST8) we diluted them to 30% (vol/vol) in TSB (Fig. 1). The toxicity of each isolate was measured using three biological repeats with high reproducibility (Suppl Fig. 1). The proportion of ST239 and ST36 isolates that killed more than 50% of the THP-1 cells was 44% and 39%, respectively. By contrast, the proportion of ST8, ST22 and ST93 isolates that killed more than 50% of cells was 87%, 84% and 85%, respectively. Given both the difference in the proportion of isolates killing more than 50% of the cells and the differences in dilutions required (100% vs 30%), this demonstrates that ST8, ST22 and ST93 contain a higher proportion of highly toxic isolates than ST239 and ST36. However, what was equally striking from this initial analysis is the scale of the variation in the toxicity of the isolates within each ST (Fig. 1).

**Figure 1:**
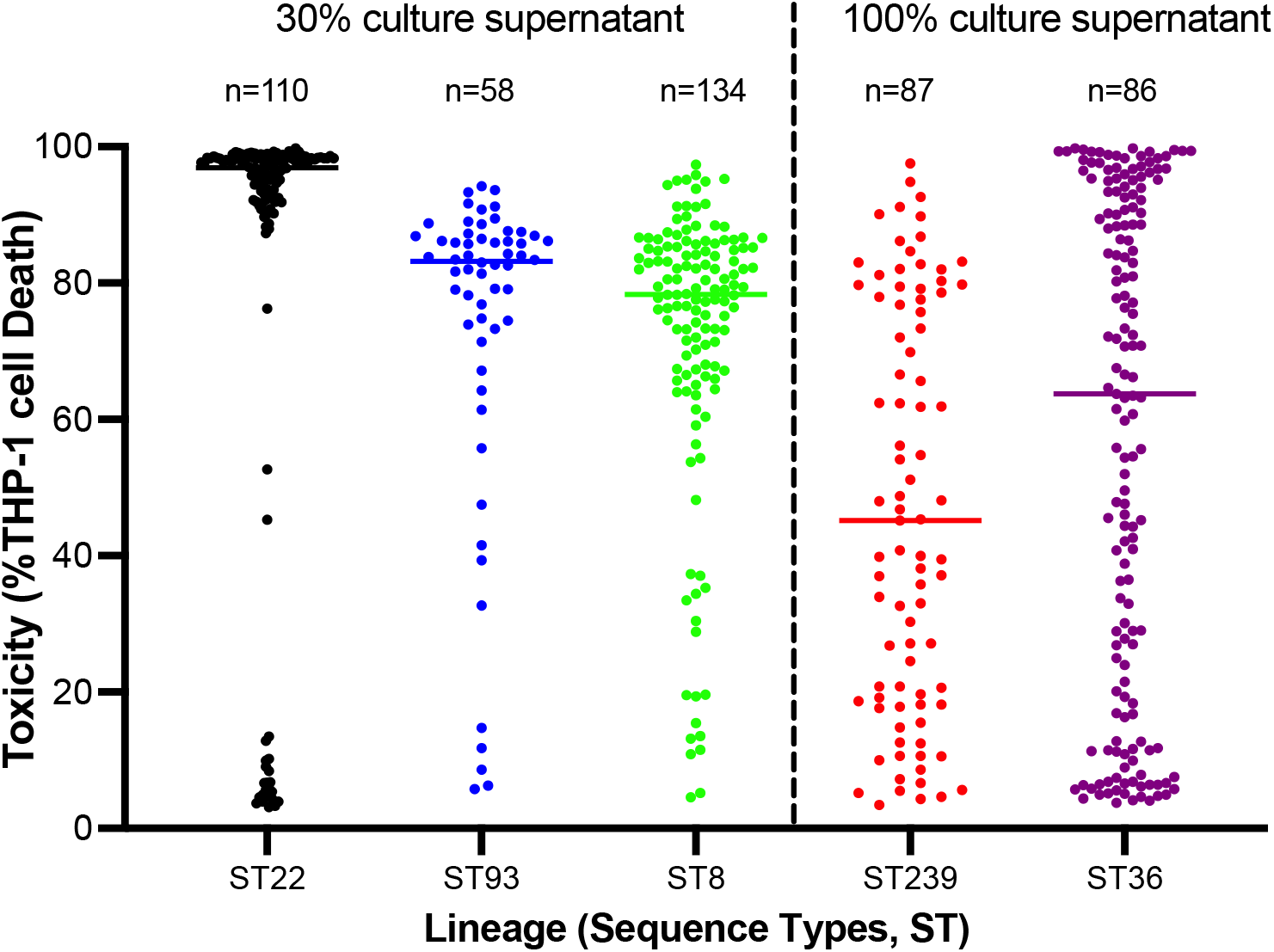
Variation in toxicity between and within MRSA sequence types. The toxicity of each isolate from five MRSA STs (ST22 (n=110), ST93 (n=58), ST8 (n=134), ST239 (n=87) and ST36 (n=86)) was quantified by incubating cell-free supernatant (either diluted to 30% using sterile TS broth or used neat (100%)) with cultured THP-1 cells and toxicity examined by flow cytometry. The toxicity of each isolate was quantified using three biological repeats with a single dot representing the mean value for each isolate and the median of each sequence type indicated by the horizontal bars.

### Toxicity cannot be inferred solely from the MRSA sequence type

To further compare the toxicity between STs we took the five most and five least toxic isolates from each MRSA ST and quantified their toxicity over a range of dilutions of their supernatant (Fig. 2). The toxicity of each supernatant dilution was measured using three biological repeats with high reproducibility (Suppl Fig. 2). The mean toxicity of the five least toxic ST8 isolates was slightly higher than those from the other STs, but this was not statistically significant (p>0.05, Fig. 2a). This demonstrates that each of the five MRSA STs contain isolates expressing comparable low levels of toxicity. Of the most toxic isolates, there were significant differences in toxicity across the five STs (Fig. 2 and Suppl Fig. 2). At a supernatant dilution of 10% the ST22 and ST8 isolates were on average more toxic than the others (*p*<0.05 for each comparison, Fig. 2b). At supernatant dilution of 30%, the ST22, ST93 and ST8 were on average more toxic than the other STs (*p*<0.05 for each comparison values), while at dilution 70%, the ST36 isolates were statistically significantly less toxic than the other STs (*p*<0.05 for each comparison values).

**Figure 2:**
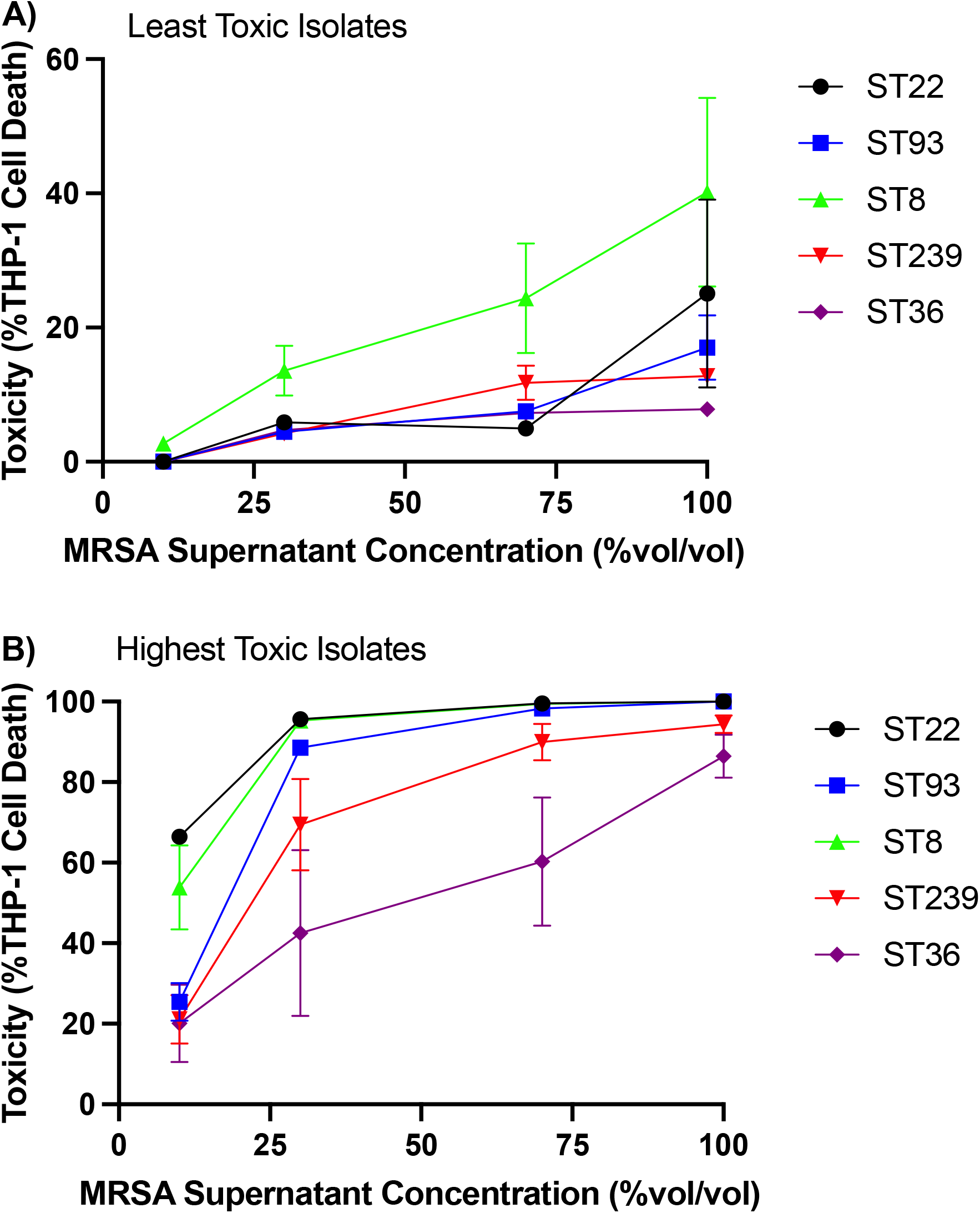
Toxicity of the least and highest toxic isolates from each of the five MRSA sequence type. The supernatant of the five least **(A)** and highest **(B)** toxic isolates from each sequence type was diluted to 10%, 30%, or 70% supernatant in sterile TS broth or used neat (100%) and the percentage cell death of THP-1 cells examined. The toxicity of each isolate was quantified using three biological repeats and the data is presented as the mean and standard deviation across the five isolates.

ST22, ST93 and ST8 all contain the community-associated Type IV SCC*mec* element, which prior to this study would have led to them being referred to as hyper-virulent (9, 25). By comparison, ST239 and ST36 isolates contain the type III and II healthcare-associated SCC*mec* elements, respectively. While the ST8 and ST22 collections did contain the most toxic isolates, the most toxic ST93 isolates were no more toxic than those from the ST239 and ST36 collections (Fig. 2B; Suppl Fig 2B). This may be a result of differential expression of toxins to which our cell line is not susceptible. However, the ST93 collection did contain a higher proportion of highly toxic isolates than the ST239 and ST36 collections, which is what we observed with the other type IV SCC*mec* carrying MRSA STs studied here. This observation aligns with earlier work demonstrating that the type IV element has less of a down-regulating effect on toxicity compared to the larger hospital associated SCC*mec* types (26).

### Agr mutations alone do not explain all low toxic isolates

To understand the molecular mechanisms behind the observed variation in toxin production, we sought to examine the impact of sequence variability within the Agr regulatory locus on this phenotype. AgrC and AgrA represent the sensor kinase and response regulator of the Agr system, respectively (Fig. 3a), and mutations within the genes encoding these key proteins are frequently associated with reduced toxin production by clinical isolates (Fig. 3a) (16). As the genome sequences were available for all the isolates studied here, we interrogated these and found that of the 475 isolates, 14 isolates had non-synonymous mutations in *agrA* (Table 1) and 35 isolates had non-synonymous mutations in *agrC* (Table 2). The location of each of the amino acid changes inferred by these mutations is indicated in Tables 1 & 2 and have been mapped onto a representation of each protein (AgrA, Fig. 3b; AgrC, Fig. 3c) where the critically active regions are indicated.

**Table 1:** Comparison of mutations identified in the accessory gene regulator A (*agrA*) gene and toxicity of MRSA isolates

| ST/Strain ID | Mutation | Description | Cell Death (%) |
| --- | --- | --- | --- |
| **ST22** |  |  |  |
| ASARM205 | Y95H | Unknown | 45 |
| ASARM93 | E163G | Glutamic residue lies within Beta-sheet 3, important in Beta-beta-beta sandwich formation, involved in salt bridge formation | 9 |
| ASARM207 | K223Stop | Dysfunctional, truncated AgrA | 5 |
| ASARM128 | K236Stop | Dysfunctional, truncated AgrA | 3 |
| **ST93** |  |  |  |
| Sa_TPS310<br>5 | Frameshift<br>I156, T178 | Dysfunctional, truncated | 6 |
| **ST8** |  |  |  |
| MR026 | (-t)2149463 | Dysfunctional, frameshift and predicted premature stop codon at 173 | 12 |
| **ST239** |  |  |  |
| AGT9 | A47D | Unknown | 6 |
| MAL119 | S139I | Functional | 91 |
| **ST30** |  |  |  |
| EOE120 | D157Y | Important in formation of elongated beta-beta-beta fold and salt bridge formation | 4 |
| ASARM63 | H169Y | Dysfunctional, His residue is essential for DNA binding | 5 |
| EOE176 | Q179Stop<br>codon | Dysfunctional, truncated AgrA | 5 |
| EOE161 | Q179Stop<br>codon | Dysfunctional, truncated AgrA | 5 |
| EOE171 | Q179Stop<br>codon | Dysfunctional, truncated AgrA | 5 |
| EOE169 | Q179Stop<br>codon | Dysfunctional, truncated AgrA | 6 |

**Table 2:** Comparison of mutations identified in the accessory gene regulator C (*agrC*) gene and toxicity of MRSA isolates

| ST/Strain ID | Mutation | Description | Cell Death (%) |
| --- | --- | --- | --- |
| **ST22** |  |  |  |
| ASARM204 | S47T & V367I | Mutation in extracellular loop; mutation in cytoplasmic c-terminal domain | 5 |
| ASARM84 | M53I | Unknown; mutation in membrane spanning region | 5 |
| ASARM224 | A57V | Functional | 91 |
| ASARM217 | A57V | Functional | 92 |
| ASARM208 | A57V | Functional | 97 |
| ASARM61 | A57V | Functional | 97 |
| ASARM200 | Y121H & Q202H | Functional | 92 |
| ASARM201 | Y121H & Q202H | Functional | 93 |
| ASARM154 | F162V | Unknown; mutation in membrane spanning region | 13 |
| ASARM97 | A340V | Mutation of highly conserved residue in N-box CA kinase subdomain | 14 |
| **ST93** |  |  |  |
| Sa_TPS3155 | Y71H | Unknown; mutation in membrane spanning region | 42 |
| Sa_TPS3165 | F162S | Unknown; mutation in membrane spanning region | 15 |
| Sa_TPS3167 | M20I | Unknown; mutation in membrane spanning region | 56 |
| Sa_TPS3148 | R235C | Mutation of conserved residue in H-box in DHp subdomain | 6 |
| Sa_TPS3151 | D240N | Mutation of conserved residue in H-box in DHp subdomain | 12 |
| Sa_TPS3161 | G284D | Mutation in histidine kinase domain - predicted to be dysfunctional | 9 |
| **ST8** |  |  |  |
| MR030 | (-t)2146811 | Deletion (-t;2146811) within intergenic region between P2 / P3 agr promoter | 64 |
| MR081 | (-t)2146811 | Deletion (-t;2146811) within intergenic region between P2 / P3 agr promoter | 23 |
| MR083 | (-t)2146811 | Deletion (-t;2146811) within intergenic region between P2 / P3 agr promoter | 65 |
| USFL093 | Y32T | Functional | 95 |
| MR065 | L381F | Mutation of non-conserved residue<br>in G-box CA kinase subdomain | 35 |
| **ST239** |  |  |  |
| GRE4 | T246M | Mutation of conserved residue in H-box in DHp subdomain | 5 |
| ICP5062 | T247I | Mutation of conserved residue in H-box in DHp subdomain | 6 |
| HU5 | I311T;<br>A343T | Dysfunctional; I311T/A343T resulted<br>in delayed RNAIII activity | 15 |
| HU11 | I311T;<br>A343T | Dysfunctional; I311T/A343T resulted<br>in delayed RNAIII activity | 18 |
| DEU29 | I311T;<br>A343T | Dysfunctional; I311T/A343T resulted<br>in delayed RNAIII activity | 19 |
| DEU37 | I311T;<br>A343T | Dysfunctional; I311T/A343T resulted<br>in delayed RNAIII activity | 19 |
| HU6 | I311T;<br>A343T | Dysfunctional; I311T/A343T resulted<br>in delayed RNAIII activity | 120 |
| DEU20 | I311T;<br>A343T | Dysfunctional; I311T/A343T resulted<br>in delayed RNAIII activity | 21 |
| DEU9 | I311T;<br>A343T | Dysfunctional; I311T/A343T resulted<br>in delayed RNAIII activity | 20 |
| DEU17 | I311T;<br>A343T | Dysfunctional; I311T/A343T resulted<br>in delayed RNAIII activity | 16 |
| HU16 | I311T;<br>A343T | Dysfunctional; I311T/A343T resulted<br>in delayed RNAIII activity | 18 |
| CHI61 | M326T | Functional | 80 |
| **ST36** |  |  |  |
| EOE173 | T247I | Mutation of conserved residue in H-box in DHp subdomain | 6 |
| EOE096 | T388I | Mutation of conserved residue in G-box CA subdomain | 7 |

**Figure 3:**
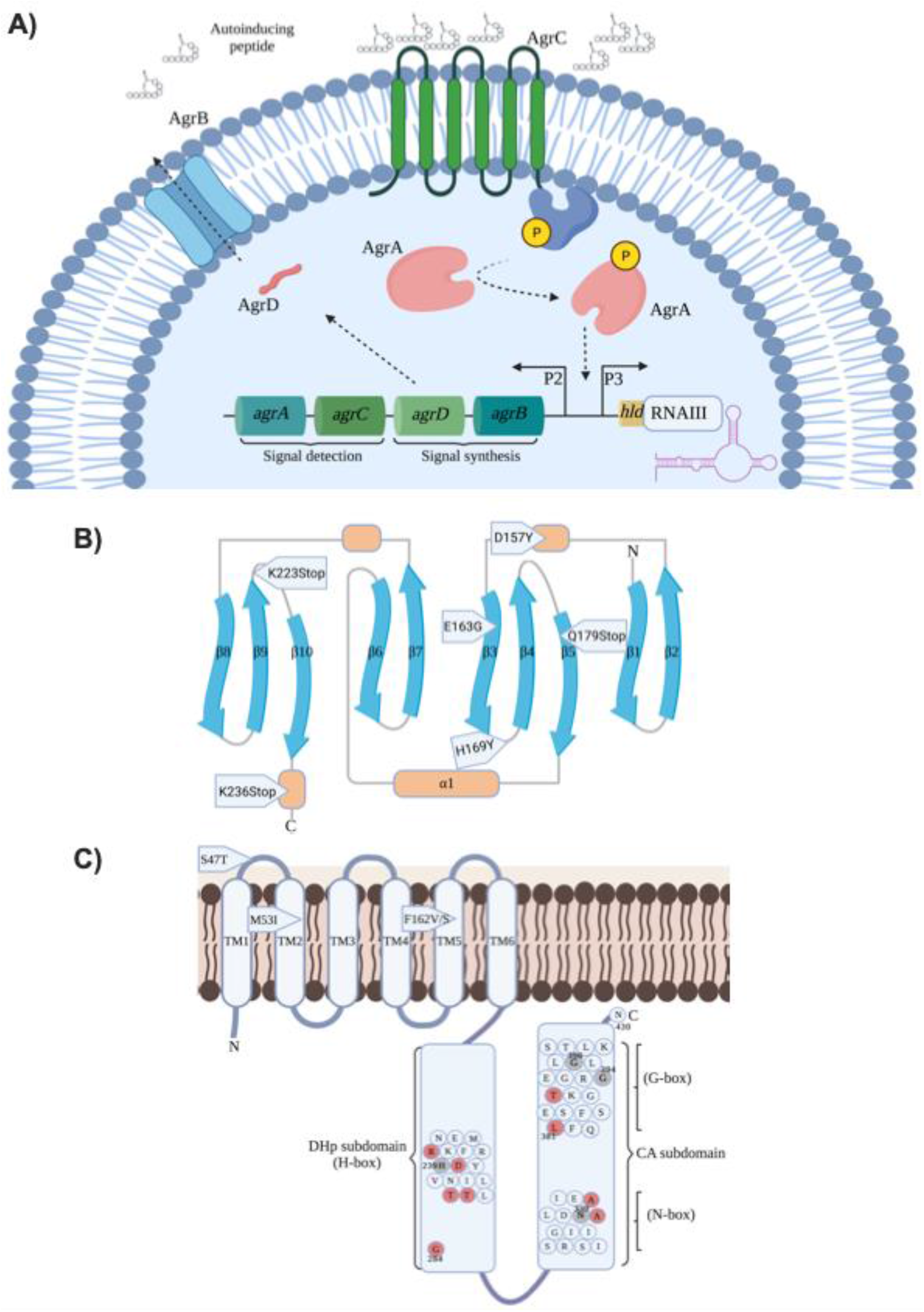
Accessory gene regulatory (Agr) system of *S aureus* labelled with mutations associated with reduced toxicity. **(A)** The Agr locus consists of two divergent transcripts driven by the P2 and P3 promoters. P2 drives the expression of the quorum sensing systems consisting of the signal synthesis (*agrBD*) and signal detection (*agrAC*) genes. AgrB and AgrD cooperative to process and secrete autoinducing peptides (AIPs) which are sensed by the polytopic transmembrane protein, AgrC. AgrCA function as a two-component signal transduction system with AgrC phosphorylating AgrA resulting in a conformational change promoting DNA binding to the intergenic region between P2 and P3 driving their expression. The effector molecule of the Agr system, RNAIII, is expressed from P3 resulting in a shift in virulence gene expression, namely enhanced cytolytic toxin expression. **B)** The C-terminal DNA binding domain of AgrA is shown as a 10-stranded elongated β-β-β sandwich, where the β-strands are shown in blue, helices shown in orange and loop regions shown in grey. Mutation associated with reduced toxicity are highlighted in the specific regions of the protein in which they occur. **C)** The transmembrane sensor and cytoplasmic histidine kinase domains of AgrC are highlighted. The central histidine residue (H239) within the H-box of the DHp subdomain and the CA subdomain N-box asparagine (N339) and glycine residues of the G box (G394 and G396) are indicated. Residues labelled in red have been identified in this study to be associated with reduced toxicity.

Given the major role the Agr system plays in toxin production of *S. aureus* we examined what proportion of low toxic isolates could be explained by these Agr mutations. We set an arbitrary threshold where we considered any isolate that killed less than 20% of the cells as ‘low toxicity’, and any that killed more than 80% as ‘high toxicity’. Of the 475 isolates, 17.7% (n=84) of the isolates were categorised as low toxicity and of these 34 (40.5%) had non-synonymous mutations in the *agr* locus. While of the high toxicity isolates (n=215 (45.2%)) eight had non-synonymous mutations in the *agr* locus, suggesting that these amino acids are not critical to the activity of the Agr system (Table 1 & 2). Of the 34 low toxicity isolates with mutations in the Agr locus, while we cannot assume these mutations are causative without making a series of isogenic mutants, having mapped them to the functional regions of the proteins (fig. 3b & c) we can infer those mutations that are likely to be causative of the low toxicity phenotype.

#### Assessment of *agrA* mutations

The AgrA protein is part of the LytR family of DNA-binding response regulators that modulate virulence determinants in several pathogenic bacteria (27). This transcription factor contains the LytTR domain which binds with high affinity to both P2 and P3 promoter regions of the *agr* locus upregulating toxin production (Fig. 3a) (16, 28). The structure of the C-terminal LytTR DNA binding domain of AgrA (residues 137-238) in complex with a DNA duplex has been solved (29), highlighting the formation of an intricate 10-stranded elongated β-β-β sandwich (Fig. 3b). Key residues within three loop regions extending from the β-sheets makes contact with the major and intervening minor groove of its DNA target resulting in increased transcription from the P2 and P3 promoters and activation of the Agr system (29). The mutations in eight of the low toxicity isolates resulted in premature truncation of the AgrA protein, which caused a loss of a functional region, and as such can be considered causative of the low toxicity phenotype (Table 1, Fig. 3b). Two of the other low toxicity isolates (ASARM93 & EOE120) had substitution mutations (E163G and D157Y, respectively) within the critical β-β-β sandwich. Mutation E163G occurs within β-strand 3 which forms the centre of the LytTR domain and plays a role in salt-bridge formation with H174 (29). Mutation D157Y is located within an α-helix between strands β2 and β3 and forms a salt bridge interaction with both H208 located on helix between β5 and β6 and E141 positioned on the beginning of strand β1. These salt bridge interactions stabilise the LytTR β-β-β fold, and our data indicates that mutations within these residues impairs Agr activity and results in reduced toxicity.

Lastly, isolate ASARM63 had a mutation conferring a H169Y change, which results in the loss of a histidine residue critical to the DNA binding activity of AgrA [32]. Only one low toxic *agrA* mutant contained a substitution in a region with no ascribed function, and that was AGT9 which had an A47D change (Not shown in Fig. 3b).

#### Assessment of *agrC* mutations

The AgrC protein is composed of a highly variable polytopic transmembrane sensor which relays auto-inducing peptide (AIP) mediated signals to the highly conserved cytoplasmic histidine kinase (HK) domain (Fig. 3a,c) (30). The AgrC HK domain is composed of two subdomains; the catalytic ATP-binding (CA) domain which promotes the autophosphorylation of the central histidine (His239) residue within the H-box of the dimerized histidine phosphotransfer (DHp) domain (Fig. 3c) (30–32). The CA subdomain N-box asparagine (N339) and glycine residues situated within the G-box (G394 and G396) are considered essential for ATP binding and Agr activity (33, 34). Importantly, alanine 340 and threonine 388 are conserved residues within this subdomain (32). Two low toxic isolates (ASARM97 (A340V) and EOE096 (T388I)) contain substitutions in the CA kinase subdomain region of AgrC, and as such may be causative of the low toxic phenotype. Six low toxic isolates (Sa_TPS3148, Sa_TPS3151, Sa_TPS3161, GRE4, ICP5062 and EOE173) occur in conserved residues within the H-box of the DHp subdomain and as such, are likely to be causative of the low toxicity phenotype. Nine isolates, all ST239 belonging to the Turkish lineage (35), contained a double substitution of I311T & A343T, and in previous work we have functionally verified that these mutations result in a significant delay to RNAIII activation (18). Molecular modelling has previously indicated that these mutations prevent AgrC dimerization and access to the ATP-binding pocket required for Agr activity which explains the low toxic phenotype (36).

Of the low toxic isolates with *agrC* mutations, four (ASARM204, ASARM84, ASARM154 and Sa_TPS3165) contain substitutions within either the extracellular or cytoplasmic regions of the protein with no ascribed function, and so we cannot claim with a level of confidence that these are causative of the low toxic phenotype.

This study adds to the growing literature indicating that toxin regulation is highly complex in *S. aureus* and confirms that there are still undiscovered mechanisms at play that modulate this major virulence phenotype. This works reinforces the importance of Agr mutations in *S. aureus* toxicity. The data generated here provides a clearer understanding of the relationship between Agr mutations and toxicity, which may be exploited for future anti-virulence drug design. However, considering that close to 60% of low toxicity isolates had no Agr mutations demonstrates that numerous regulatory mechanisms await discovery, and work dedicated to unravelling the regulatory circuits controlling the toxicity of *S. aureus* is ongoing. This also suggests that use of the term ‘Agr dysfunction’ should be used with consideration of the fact that for many of low toxicity clinical isolates the Agr system is likely to be functional.

What this work primarily highlights is the care and consideration needed when inferring terms like ‘hypervirulence’ to an isolate based on its sequence type. There can be no doubt that CA-MRSA lineages have recently emerged and are highly successful. Whether this is a result of hyper-virulence’ as opposed to an enhanced ability to transmit amongst otherwise healthy individuals needs further investigation. From the perspective of cytolytic toxin production by MRSA, the level of variability is significant and likely to play a major role in the outcome of *S. aureus disease.*

## Author contributions

M.L. and R.C.M. conceived the study and wrote the original manuscript, M.L., B.B., and S.L.B. performed experiments and analysed data, S.J.P., B.B., S.L.B., B.J.H., and T.P.S. participated in writing the manuscript and supported the project.

## Abbreviations

(ST): Sequence type
(MRSA): methicillin-resistant *S. aureus*
(CA-MRSA): community-associated MRSA
(Agr): accessory gene regulator
(TSB): Tryptic-Soy Broth
(HK): histidine kinase
(CA domain): catalytic ATP-binding domain
(DHp domain): histidine phosphotransfer domain

## References

1. Gordon RJ, Lowy FD. Pathogenesis of methicillin-resistant Staphylococcus aureus infection. Clinical infectious diseases : an official publication of the Infectious Diseases Society of America. 2008;46 Suppl 5:S350–9.

2. Chambers HF, Deleo FR. Waves of resistance: Staphylococcus aureus in the antibiotic era. Nature reviews Microbiology. 2009;7(9):629–41.

3. Chambers HF. The changing epidemiology of Staphylococcus aureus? Emerging infectious diseases. 2001;7(2):178–82.

4. David MZ, Daum RS. Community-associated methicillin-resistant Staphylococcus aureus: epidemiology and clinical consequences of an emerging epidemic. Clin Microbiol Rev. 2010;23(3):616–87.

5. Spaan AN, Henry T, van Rooijen WJM, Perret M, Badiou C, Aerts PC, et al. The staphylococcal toxin Panton-Valentine Leukocidin targets human C5a receptors. Cell Host Microbe. 2013;13(5):584–94.

6. Diep BA, Chan L, Tattevin P, Kajikawa O, Martin TR, Basuino L, et al. Polymorphonuclear leukocytes mediate Staphylococcus aureus Panton-Valentine leukocidin-induced lung inflammation and injury. Proc Natl Acad Sci U S A. 2010;107(12):5587–92.

7. Kobayashi SD, Malachowa N, Whitney AR, Braughton KR, Gardner DJ, Long D, et al. Comparative analysis of USA300 virulence determinants in a rabbit model of skin and soft tissue infection. J Infect Dis. 2011;204(6):937–41.

8. Lipinska U, Hermans K, Meulemans L, Dumitrescu O, Badiou C, Duchateau L, et al. Panton-Valentine leukocidin does play a role in the early stage of Staphylococcus aureus skin infections: a rabbit model. PLoS One. 2011;6(8):e22864.

9. Li M, Diep BA, Villaruz AE, Braughton KR, Jiang X, DeLeo FR, et al. Evolution of virulence in epidemic community-associated methicillin-resistant Staphylococcus aureus. Proc Natl Acad Sci U S A. 2009;106(14):5883–8.

10. Li M, Cheung GY, Hu J, Wang D, Joo HS, Deleo FR, et al. Comparative analysis of virulence and toxin expression of global community-associated methicillin-resistant Staphylococcus aureus strains. J Infect Dis. 2010;202(12):1866–76.

11. Wang R, Braughton KR, Kretschmer D, Bach TH, Queck SY, Li M, et al. Identification of novel cytolytic peptides as key virulence determinants for community-associated MRSA. Nat Med. 2007;13(12):1510–4.

12. Chua KY, Monk IR, Lin YH, Seemann T, Tuck KL, Porter JL, et al. Hyperexpression of alpha-hemolysin explains enhanced virulence of sequence type 93 community-associated methicillin-resistant Staphylococcus aureus. BMC Microbiol. 2014;14:31.

13. Song L, Hobaugh MR, Shustak C, Cheley S, Bayley H, Gouaux JE. Structure of staphylococcal alpha-hemolysin, a heptameric transmembrane pore. Science. 1996;274(5294):1859–66.

14. Berube BJ, Bubeck Wardenburg J. Staphylococcus aureus alpha-toxin: nearly a century of intrigue. Toxins (Basel). 2013;5(6):1140–66.

15. Laabei M, Jamieson WD, Yang Y, van den Elsen J, Jenkins AT. Investigating the lytic activity and structural properties of Staphylococcus aureus phenol soluble modulin (PSM) peptide toxins. Biochim Biophys Acta. 2014;1838(12):3153–61.

16. Novick RP. Autoinduction and signal transduction in the regulation of staphylococcal virulence. Mol Microbiol. 2003;48(6):1429–49.

17. Cheung GY, Wang R, Khan BA, Sturdevant DE, Otto M. Role of the accessory gene regulator agr in community-associated methicillin-resistant Staphylococcus aureus pathogenesis. Infect Immun. 2011;79(5):1927–35.

18. Laabei M, Recker M, Rudkin JK, Aldeljawi M, Gulay Z, Sloan TJ, et al. Predicting the virulence of MRSA from its genome sequence. Genome Res. 2014;24(5):839–49.

19. Wilke GA, Bubeck Wardenburg J. Role of a disintegrin and metalloprotease 10 in Staphylococcus aureus alpha-hemolysin-mediated cellular injury. Proc Natl Acad Sci U S A. 2010;107(30):13473–8.

20. Laabei M, Uhlemann AC, Lowy FD, Austin ED, Yokoyama M, Ouadi K, et al. Evolutionary Trade-Offs Underlie the Multi-faceted Virulence of Staphylococcus aureus. PLoS Biol. 2015;13(9):e1002229.

21. Chua K, Seemann T, Harrison PF, Davies JK, Coutts SJ, Chen H, et al. Complete genome sequence of Staphylococcus aureus strain JKD6159, a unique Australian clone of ST93-IV community methicillin-resistant Staphylococcus aureus. J Bacteriol. 2010;192(20):5556–7.

22. Chua KY, Seemann T, Harrison PF, Monagle S, Korman TM, Johnson PD, et al. The dominant Australian community-acquired methicillin-resistant Staphylococcus aureus clone ST93-IV [2B] is highly virulent and genetically distinct. PLoS One. 2011;6(10):e25887.

23. Harris SR, Feil EJ, Holden MT, Quail MA, Nickerson EK, Chantratita N, et al. Evolution of MRSA during hospital transmission and intercontinental spread. Science. 2010;327(5964):469–74.

24. Recker M, Laabei M, Toleman MS, Reuter S, Saunderson RB, Blane B, et al. Clonal differences in Staphylococcus aureus bacteraemia-associated mortality. Nat Microbiol. 2017;2(10):1381–8.

25. Otto M. Basis of virulence in community-associated methicillin-resistant Staphylococcus aureus. Annu Rev Microbiol. 2010;64:143–62.

26. Rudkin JK, Edwards AM, Bowden MG, Brown EL, Pozzi C, Waters EM, et al. Methicillin resistance reduces the virulence of healthcare-associated methicillin-resistant Staphylococcus aureus by interfering with the agr quorum sensing system. The Journal of infectious diseases. 2012;205(5):798–806.

27. Nikolskaya AN, Galperin MY. A novel type of conserved DNA-binding domain in the transcriptional regulators of the AlgR/AgrA/LytR family. Nucleic Acids Res. 2002;30(11):2453–9.

28. Koenig RL, Ray JL, Maleki SJ, Smeltzer MS, Hurlburt BK. Staphylococcus aureus AgrA binding to the RNAIII-agr regulatory region. J Bacteriol. 2004;186(22):7549–55.

29. Sidote DJ, Barbieri CM, Wu T, Stock AM. Structure of the Staphylococcus aureus AgrA LytTR domain bound to DNA reveals a beta fold with an unusual mode of binding. Structure. 2008;16(5):727–35.

30. Grebe TW, Stock JB. The histidine protein kinase superfamily. Adv Microb Physiol. 1999;41:139–227.

31. Geisinger E, Muir TW, Novick RP. agr receptor mutants reveal distinct modes of inhibition by staphylococcal autoinducing peptides. Proc Natl Acad Sci U S A. 2009;106(4):1216–21.

32. George Cisar EA, Geisinger E, Muir TW, Novick RP. Symmetric signalling within asymmetric dimers of the Staphylococcus aureus receptor histidine kinase AgrC. Mol Microbiol. 2009;74(1):44–57.

33. Hirschman A, Boukhvalova M, VanBruggen R, Wolfe AJ, Stewart RC. Active site mutations in CheA, the signal-transducing protein kinase of the chemotaxis system in Escherichia coli. Biochemistry. 2001;40(46):13876–87.

34. Zhu Y, Inouye M. The role of the G2 box, a conserved motif in the histidine kinase superfamily, in modulating the function of EnvZ. Mol Microbiol. 2002;45(3):653–63.

35. Castillo-Ramirez S, Corander J, Marttinen P, Aldeljawi M, Hanage WP, Westh H, et al. Phylogeographic variation in recombination rates within a global clone of methicillin-resistant Staphylococcus aureus. Genome Biol. 2012;13(12):R126.

36. Sloan TJ, Murray E, Yokoyama M, Massey RC, Chan WC, Bonev BB, et al. Timing Is Everything: Impact of Naturally Occurring Staphylococcus aureus AgrC Cytoplasmic Domain Adaptive Mutations on Autoinduction. J Bacteriol. 2019;201(20).

