## Supplementary material for "Significant Variability exists in the Toxicity of Global Methicillin-resistant *Staphylococcus aureus* Lineages": Supp figures and data

**Supplementary Table 1:** List of clinical isolates used in this study

| Clinical Isolate | Sequence Type | Country of Origin | Disease Status | Year |
|---|---|---|---|---|
| ASARM100 | ST22 | UK | Bacteraemia | 2007 |
| ASARM101 | ST22 | UK | Bacteraemia | 2007 |
| ASARM102 | ST22 | UK | Bacteraemia | 2007 |
| ASARM103 | ST22 | UK | Bacteraemia | 2007 |
| ASARM105 | ST22 | UK | Bacteraemia | 2007 |
| ASARM107 | ST22 | UK | Bacteraemia | 2007 |
| ASARM108 | ST22 | UK | Bacteraemia | 2007 |
| ASARM109 | ST22 | UK | Bacteraemia | 2007 |
| ASARM110 | ST22 | UK | Bacteraemia | 2007 |
| ASARM114 | ST22 | UK | Bacteraemia | 2007 |
| ASARM116 | ST22 | UK | Bacteraemia | 2007 |
| ASARM117 | ST22 | UK | Bacteraemia | 2007 |
| ASARM118 | ST22 | UK | Bacteraemia | 2007 |
| ASARM119 | ST22 | UK | Bacteraemia | 2007 |
| ASARM120 | ST22 | UK | Bacteraemia | 2007 |
| ASARM121 | ST22 | UK | Bacteraemia | 2007 |
| ASARM122 | ST22 | UK | Bacteraemia | 2007 |
| ASARM124 | ST22 | UK | Bacteraemia | 2007 |
| ASARM125 | ST22 | UK | Bacteraemia | 2007 |
| ASARM126 | ST22 | UK | Bacteraemia | 2007 |
| ASARM127 | ST22 | UK | Bacteraemia | 2007 |
| ASARM128 | ST22 | UK | Bacteraemia | 2007 |
| ASARM132 | ST22 | UK | Bacteraemia | 2007 |
| ASARM133 | ST22 | UK | Bacteraemia | 2007 |
| ASARM134 | ST22 | UK | Bacteraemia | 2007 |
| ASARM135 | ST22 | UK | Bacteraemia | 2007 |
| ASARM136 | ST22 | UK | Bacteraemia | 2007 |
| ASARM137 | ST22 | UK | Bacteraemia | 2007 |
| ASARM138 | ST22 | UK | Bacteraemia | 2007 |
| ASARM139 | ST22 | UK | Bacteraemia | 2007 |
| ASARM140 | ST22 | UK | Bacteraemia | 2007 |
| ASARM141 | ST22 | UK | Bacteraemia | 2007 |
| ASARM142 | ST22 | UK | Bacteraemia | 2007 |
| ASARM143 | ST22 | UK | Bacteraemia | 2007 |
| ASARM144 | ST22 | UK | Bacteraemia | 2007 |
| ASARM145 | ST22 | UK | Bacteraemia | 2007 |
| ASARM148 | ST22 | UK | Bacteraemia | 2008 |
| ASARM153 | ST22 | UK | Bacteraemia | 2008 |
| ASARM154 | ST22 | UK | Bacteraemia | 2008 |
| ASARM155 | ST22 | UK | Bacteraemia | 2008 |

| | | | | |
|---|---|---|---|---|
| ASARM160 | ST22 | UK | Bacteraemia | 2008 |
| ASARM162 | ST22 | UK | Bacteraemia | 2008 |
| ASARM163 | ST22 | UK | Bacteraemia | 2008 |
| ASARM164 | ST22 | UK | Bacteraemia | 2008 |
| ASARM165 | ST22 | UK | Bacteraemia | 2008 |
| ASARM166 | ST22 | UK | Bacteraemia | 2008 |
| ASARM167 | ST22 | UK | Bacteraemia | 2008 |
| ASARM168 | ST22 | UK | Bacteraemia | 2008 |
| ASARM169 | ST22 | UK | Bacteraemia | 2008 |
| ASARM170 | ST22 | UK | Bacteraemia | 2008 |
| ASARM171 | ST22 | UK | Bacteraemia | 2008 |
| ASARM172 | ST22 | UK | Bacteraemia | 2008 |
| ASARM176 | ST22 | UK | Bacteraemia | 2009 |
| ASARM177 | ST22 | UK | Bacteraemia | 2009 |
| ASARM179 | ST22 | UK | Bacteraemia | 2009 |
| ASARM181 | ST22 | UK | Bacteraemia | 2009 |
| ASARM183 | ST22 | UK | Bacteraemia | 2009 |
| ASARM184 | ST22 | UK | Bacteraemia | 2009 |
| ASARM191 | ST22 | UK | Bacteraemia | 2009 |
| ASARM193 | ST22 | UK | Bacteraemia | 2009 |
| ASARM195 | ST22 | UK | Bacteraemia | 2009 |
| ASARM196 | ST22 | UK | Bacteraemia | 2009 |
| ASARM199 | ST22 | UK | Bacteraemia | 2009 |
| ASARM200 | ST22 | UK | Bacteraemia | 2010 |
| ASARM201 | ST22 | UK | Bacteraemia | 2010 |
| ASARM203 | ST22 | UK | Bacteraemia | 2010 |
| ASARM204 | ST22 | UK | Bacteraemia | 2010 |
| ASARM205 | ST22 | UK | Bacteraemia | 2010 |
| ASARM207 | ST22 | UK | Bacteraemia | 2010 |
| ASARM208 | ST22 | UK | Bacteraemia | 2010 |
| ASARM209 | ST22 | UK | Bacteraemia | 2010 |
| ASARM211 | ST22 | UK | Bacteraemia | 2010 |
| ASARM212 | ST22 | UK | Bacteraemia | 2010 |
| ASARM217 | ST22 | UK | Bacteraemia | 2011 |
| ASARM220 | ST22 | UK | Bacteraemia | 2011 |
| ASARM221 | ST22 | UK | Bacteraemia | 2011 |
| ASARM222 | ST22 | UK | Bacteraemia | 2012 |
| ASARM223 | ST22 | UK | Bacteraemia | 2012 |
| ASARM224 | ST22 | UK | Bacteraemia | 2012 |
| ASARM59 | ST22 | UK | Bacteraemia | 2006 |
| ASARM61 | ST22 | UK | Bacteraemia | 2006 |
| ASARM62 | ST22 | UK | Bacteraemia | 2006 |
| ASARM64 | ST22 | UK | Bacteraemia | 2006 |

| | | | | |
|---|---|---|---|---|
| ASARM65 | ST22 | UK | Bacteraemia | 2006 |
| ASARM67 | ST22 | UK | Bacteraemia | 2006 |
| ASARM68 | ST22 | UK | Bacteraemia | 2006 |
| ASARM69 | ST22 | UK | Bacteraemia | 2006 |
| ASARM70 | ST22 | UK | Bacteraemia | 2006 |
| ASARM71 | ST22 | UK | Bacteraemia | 2006 |
| ASARM72 | ST22 | UK | Bacteraemia | 2006 |
| ASARM73 | ST22 | UK | Bacteraemia | 2006 |
| ASARM74 | ST22 | UK | Bacteraemia | 2006 |
| ASARM75 | ST22 | UK | Bacteraemia | 2006 |
| ASARM76 | ST22 | UK | Bacteraemia | 2006 |
| ASARM77 | ST22 | UK | Bacteraemia | 2006 |
| ASARM79 | ST22 | UK | Bacteraemia | 2006 |
| ASARM80 | ST22 | UK | Bacteraemia | 2006 |
| ASARM83 | ST22 | UK | Bacteraemia | 2006 |
| ASARM84 | ST22 | UK | Bacteraemia | 2006 |
| ASARM86 | ST22 | UK | Bacteraemia | 2006 |
| ASARM87 | ST22 | UK | Bacteraemia | 2006 |
| ASARM89 | ST22 | UK | Bacteraemia | 2006 |
| ASARM93 | ST22 | UK | Bacteraemia | 2006 |
| ASARM95 | ST22 | UK | Bacteraemia | 2006 |
| ASARM96 | ST22 | UK | Bacteraemia | 2006 |
| ASARM97 | ST22 | UK | Bacteraemia | 2006 |
| ASARM99 | ST22 | UK | Bacteraemia | 2007 |
| ASARMLT1 | ST22 | UK | Bacteraemia | 2008 |
| ASARMLT2 | ST22 | UK | Bacteraemia | 2008 |
| ASARMLT3 | ST22 | UK | Bacteraemia | 2008 |
| Sa_TPS3105 | ST93 | Australia | Bacteraemia | 2005 |
| Sa_TPS3148 | ST93 | Australia | Bacteraemia | 2007 |
| Sa_TPS3161 | ST93 | Australia | SSTI | 2008 |
| Sa_TPS3151 | ST93 | Australia | SSTI | 2008 |
| Sa_TPS3165 | ST93 | Australia | Carriage | 2008 |
| Sa_TPS3150 | ST93 | Australia | Sputum | 2007 |
| Sa_TPS3171 | ST93 | Australia | Carriage | 2008 |
| Sa_TPS3155 | ST93 | Australia | Carriage | 2008 |
| Sa_TPS3118 | ST93 | Australia | not recorded | not recorded |
| Sa_TPS3167 | ST93 | Australia | endotrachael tube | 2008 |
| Sa_TPS3162 | ST93 | Australia | Carriage | 2003 |
| Sa_TPS3169 | ST93 | Australia | SSTI | 2008 |
| Sa_TPS3106 | ST93 | Australia | Carriage | 2008 |
| Sa_TPS3145 | ST93 | Australia | SSTI | 2006 |
| Sa_TPS3158 | ST93 | Australia | SSTI | 2008 |
| Sa_TPS3183 | ST93 | Australia | not recorded | 2007 |
| Sa_TPS3134 | ST93 | Australia | SSTI | 2000 |

| | | | | |
|---|---|---|---|---|
| Sa_TPS3137 | ST93 | Australia | SSTI | 2000 |
| Sa_TPS3181 | ST93 | Australia | SSTI | 2008 |
| Sa_TPS3026 | ST93 | Australia | Bacteraemia | 2004 |
| Sa_TPS3142 | ST93 | Australia | Bacteraemia | 2005 |
| Sa_TPS3164 | ST93 | Australia | Carriage | 2008 |
| Sa_TPS3153 | ST93 | Australia | Carriage | 2008 |
| Sa_TPS3104 | ST93 | Australia | Carriage | 2009 |
| Sa_TPS3139 | ST93 | Australia | SSTI | 2000 |
| Sa_TPS3149 | ST93 | Australia | Carriage | 2007 |
| Sa_TPS3138 | ST93 | Australia | SSTI | 2000 |
| Sa_TPS3160 | ST93 | Australia | SSTI | 2008 |
| Sa_TPS3185 | ST93 | Australia | Carriage | 2003 |
| Sa_TPS3176 | ST93 | Australia | SSTI | 2009 |
| Sa_TPS3144 | ST93 | Australia | SSTI | 2006 |
| Sa_TPS3174 | ST93 | Australia | SSTI | 2009 |
| Sa_TPS3146 | ST93 | Australia | SSTI | 2006 |
| Sa_TPS3133 | ST93 | Australia | Carriage | 2000 |
| Sa_TPS3132 | ST93 | Australia | not recorded | 2000 |
| Sa_TPS3152 | ST93 | Australia | pneumoniae | 2008 |
| Sa_TPS3135 | ST93 | Australia | Carriage | 2000 |
| Sa_TPS3173 | ST93 | Australia | Carriage | 2009 |
| Sa_TPS3140 | ST93 | Australia | SSTI | 2000 |
| Sa_TPS3136 | ST93 | Australia | Bacteraemia | 2000 |
| Sa_TPS3188 | ST93 | Australia | not recorded | 1992 |
| Sa_TPS3166 | ST93 | Australia | Carriage | 2008 |
| Sa_TPS3178 | ST93 | Australia | Pleural fluid | 2008 |
| Sa_TPS3157 | ST93 | Australia | SSTI | 2008 |
| Sa_TPS3168 | ST93 | Australia | SSTI | 2008 |
| Sa_TPS3189 | ST93 | Australia | not recorded | 1992 |
| Sa_TPS3154 | ST93 | Australia | SSTI | 2008 |
| Sa_TPS3163 | ST93 | Australia | SSTI | 2005 |
| Sa_TPS3184 | ST93 | Australia | Carriage | 1995 |
| Sa_TPS3182 | ST93 | Australia | not recorded | 2008 |
| Sa_TPS3147 | ST93 | Australia | SSTI | 2006 |
| Sa_TPS3177 | ST93 | Australia | Bacteraemia | 2008 |
| Sa_TPS3159 | ST93 | Australia | SSTI | 2008 |
| Sa_TPS3156 | ST93 | Australia | Carriage | 2008 |
| Sa_TPS3180 | ST93 | Australia | not recorded | not recorded |
| Sa_TPS3186 | ST93 | Australia | Carriage | 2003 |
| Sa_TPS3179 | ST93 | Australia | Carriage | 2008 |
| Sa_TPS3187 | ST93 | Australia | Carriage | 1996 |
| MR005 | ST8 | USA | Bacteraemia | 2009-2011 |
| MR007 | ST8 | USA | Bacteraemia | 2009-2011 |
| MR018 | ST8 | USA | Bacteraemia | 2009-2011 |
| MR019 | ST8 | USA | Bacteraemia | 2009-2011 |
| MR021 | ST8 | USA | Bacteraemia | 2009-2011 |

| | | | | |
|---|---|---|---|---|
| MR022 | ST8 | USA | Bacteraemia | 2009-2011 |
| MR023 | ST8 | USA | Bacteraemia | 2009-2011 |
| MR025 | ST8 | USA | Bacteraemia | 2009-2011 |
| MR026 | ST8 | USA | Bacteraemia | 2009-2011 |
| MR027 | ST8 | USA | Bacteraemia | 2009-2011 |
| MR029 | ST8 | USA | Bacteraemia | 2009-2011 |
| MR030 | ST8 | USA | Bacteraemia | 2009-2011 |
| MR031 | ST8 | USA | Bacteraemia | 2009-2011 |
| MR035 | ST8 | USA | Bacteraemia | 2009-2011 |
| MR036 | ST8 | USA | Bacteraemia | 2009-2011 |
| MR039 | ST8 | USA | Bacteraemia | 2009-2011 |
| MR047 | ST8 | USA | Bacteraemia | 2009-2011 |
| MR051 | ST8 | USA | Bacteraemia | 2009-2011 |
| MR060 | ST8 | USA | Bacteraemia | 2009-2011 |
| MR063 | ST8 | USA | Bacteraemia | 2009-2011 |
| MR064 | ST8 | USA | Bacteraemia | 2009-2011 |
| MR065 | ST8 | USA | Bacteraemia | 2009-2011 |
| MR072 | ST8 | USA | Bacteraemia | 2009-2011 |
| MR073 | ST8 | USA | Bacteraemia | 2009-2011 |
| MR074 | ST8 | USA | Bacteraemia | 2009-2011 |
| MR077 | ST8 | USA | Bacteraemia | 2009-2011 |
| MR078 | ST8 | USA | Bacteraemia | 2009-2011 |
| MR081 | ST8 | USA | Bacteraemia | 2009-2011 |
| MR083 | ST8 | USA | Bacteraemia | 2009-2011 |
| MR084 | ST8 | USA | Bacteraemia | 2009-2011 |
| MR087 | ST8 | USA | Bacteraemia | 2009-2011 |
| MR090 | ST8 | USA | Bacteraemia | 2009-2011 |
| MR091 | ST8 | USA | Bacteraemia | 2009-2011 |
| MR096 | ST8 | USA | Bacteraemia | 2009-2011 |
| MR107 | ST8 | USA | Bacteraemia | 2009-2011 |
| MR110 | ST8 | USA | Bacteraemia | 2009-2011 |
| USFL008 | ST8 | USA | Carriage | 2009-2011 |
| USFL009 | ST8 | USA | Carriage | 2009-2011 |
| USFL012 | ST8 | USA | Carriage | 2009-2011 |
| USFL028 | ST8 | USA | Carriage | 2009-2011 |
| USFL042 | ST8 | USA | Carriage | 2009-2011 |
| USFL061 | ST8 | USA | Carriage | 2009-2011 |
| USFL063 | ST8 | USA | Carriage | 2009-2011 |
| USFL074 | ST8 | USA | Carriage | 2009-2011 |
| USFL077 | ST8 | USA | Carriage | 2009-2011 |
| USFL082 | ST8 | USA | Carriage | 2009-2011 |
| USFL093 | ST8 | USA | Carriage | 2009-2011 |
| USFL119 | ST8 | USA | Carriage | 2009-2011 |
| USFL130 | ST8 | USA | Carriage | 2009-2011 |
| USFL141 | ST8 | USA | Carriage | 2009-2011 |
| USFL153 | ST8 | USA | Carriage | 2009-2011 |

| | | | | |
|---|---|---|---|---|
| USFL156 | ST8 | USA | Carriage | 2009-2011 |
| USFL166 | ST8 | USA | Carriage | 2009-2011 |
| USFL167 | ST8 | USA | Carriage | 2009-2011 |
| USFL169 | ST8 | USA | Carriage | 2009-2011 |
| USFL182 | ST8 | USA | Carriage | 2009-2011 |
| USFL200 | ST8 | USA | Carriage | 2009-2011 |
| USFL211 | ST8 | USA | Carriage | 2009-2011 |
| USFL213 | ST8 | USA | Carriage | 2009-2011 |
| USFL224 | ST8 | USA | Carriage | 2009-2011 |
| USFL225 | ST8 | USA | Carriage | 2009-2011 |
| USFL230 | ST8 | USA | Carriage | 2009-2011 |
| USFL231 | ST8 | USA | Carriage | 2009-2011 |
| USFL243 | ST8 | USA | Carriage | 2009-2011 |
| USFL248 | ST8 | USA | Carriage | 2009-2011 |
| USFL259 | ST8 | USA | Carriage | 2009-2011 |
| USFL263 | ST8 | USA | Carriage | 2009-2011 |
| USFL267 | ST8 | USA | Carriage | 2009-2011 |
| USFL269 | ST8 | USA | Carriage | 2009-2011 |
| USFL271 | ST8 | USA | Carriage | 2009-2011 |
| USFL272 | ST8 | USA | Carriage | 2009-2011 |
| USFL302 | ST8 | USA | Carriage | 2009-2011 |
| USFL303 | ST8 | USA | Carriage | 2009-2011 |
| USFL304 | ST8 | USA | Carriage | 2009-2011 |
| USFL016 | ST8 | USA | SSTI | 2009-2011 |
| USFL018 | ST8 | USA | SSTI | 2009-2011 |
| USFL020 | ST8 | USA | SSTI | 2009-2011 |
| USFL021 | ST8 | USA | SSTI | 2009-2011 |
| USFL034 | ST8 | USA | SSTI | 2009-2011 |
| USFL035 | ST8 | USA | SSTI | 2009-2011 |
| USFL036 | ST8 | USA | SSTI | 2009-2011 |
| USFL039 | ST8 | USA | SSTI | 2009-2011 |
| USFL056 | ST8 | USA | SSTI | 2009-2011 |
| USFL057 | ST8 | USA | SSTI | 2009-2011 |
| USFL059 | ST8 | USA | SSTI | 2009-2011 |
| USFL069 | ST8 | USA | SSTI | 2009-2011 |
| USFL095 | ST8 | USA | SSTI | 2009-2011 |
| USFL097 | ST8 | USA | SSTI | 2009-2011 |
| USFL103 | ST8 | USA | SSTI | 2009-2011 |
| USFL110 | ST8 | USA | SSTI | 2009-2011 |
| USFL111 | ST8 | USA | SSTI | 2009-2011 |
| USFL113 | ST8 | USA | SSTI | 2009-2011 |
| USFL136 | ST8 | USA | SSTI | 2009-2011 |
| USFL137 | ST8 | USA | SSTI | 2009-2011 |
| USFL138 | ST8 | USA | SSTI | 2009-2011 |
| USFL139 | ST8 | USA | SSTI | 2009-2011 |
| USFL149 | ST8 | USA | SSTI | 2009-2011 |

| | | | | |
|---|---|---|---|---|
| USFL152 | ST8 | USA | SSTI | 2009-2011 |
| USFL158 | ST8 | USA | SSTI | 2009-2011 |
| USFL159 | ST8 | USA | SSTI | 2009-2011 |
| USFL160 | ST8 | USA | SSTI | 2009-2011 |
| USFL162 | ST8 | USA | SSTI | 2009-2011 |
| USFL164 | ST8 | USA | SSTI | 2009-2011 |
| USFL165 | ST8 | USA | SSTI | 2009-2011 |
| USFL173 | ST8 | USA | SSTI | 2009-2011 |
| USFL174 | ST8 | USA | SSTI | 2009-2011 |
| USFL194 | ST8 | USA | SSTI | 2009-2011 |
| USFL198 | ST8 | USA | SSTI | 2009-2011 |
| USFL218 | ST8 | USA | SSTI | 2009-2011 |
| USFL219 | ST8 | USA | SSTI | 2009-2011 |
| USFL220 | ST8 | USA | SSTI | 2009-2011 |
| USFL221 | ST8 | USA | SSTI | 2009-2011 |
| USFL222 | ST8 | USA | SSTI | 2009-2011 |
| USFL223 | ST8 | USA | SSTI | 2009-2011 |
| USFL237 | ST8 | USA | SSTI | 2009-2011 |
| USFL239 | ST8 | USA | SSTI | 2009-2011 |
| USFL240 | ST8 | USA | SSTI | 2009-2011 |
| USFL250 | ST8 | USA | SSTI | 2009-2011 |
| USFL255 | ST8 | USA | SSTI | 2009-2011 |
| USFL256 | ST8 | USA | SSTI | 2009-2011 |
| USFL258 | ST8 | USA | SSTI | 2009-2011 |
| USFL273 | ST8 | USA | SSTI | 2009-2011 |
| USFL274 | ST8 | USA | SSTI | 2009-2011 |
| USFL276 | ST8 | USA | SSTI | 2009-2011 |
| USFL277 | ST8 | USA | SSTI | 2009-2011 |
| USFL279 | ST8 | USA | SSTI | 2009-2011 |
| USFL282 | ST8 | USA | SSTI | 2009-2011 |
| USFL319 | ST8 | USA | SSTI | 2009-2011 |
| USFL320 | ST8 | USA | SSTI | 2009-2011 |
| USFL326 | ST8 | USA | SSTI | 2009-2011 |
| USFL327 | ST8 | USA | SSTI | 2009-2011 |
| USFL330 | ST8 | USA | SSTI | 2009-2011 |
| USFL339 | ST8 | USA | SSTI | 2009-2011 |
| USFL341 | ST8 | USA | SSTI | 2009-2011 |
| ASARM112 | ST36 | UK | Bacteraemia | 2003 |
| ASARM156 | ST36 | UK | Bacteraemia | 2004 |
| ASARM161 | ST36 | UK | Bacteraemia | 2004 |
| ASARM180 | ST36 | UK | Bacteraemia | 2005 |
| ASARM185 | ST36 | UK | Bacteraemia | 2005 |
| ASARM190 | ST36 | UK | Bacteraemia | 2005 |
| ASARM197 | ST36 | UK | Bacteraemia | 2005 |
| ASARM210 | ST36 | UK | Bacteraemia | 2006 |
| ASARM213 | ST36 | UK | Bacteraemia | 2006 |

| | | | | |
|---|---|---|---|---|
| ASARM63 | ST36 | UK | Bacteraemia | 2002 |
| ASARM88 | ST36 | UK | Bacteraemia | 2002 |
| ASARM90 | ST36 | UK | Bacteraemia | 2002 |
| ASARM92 | ST36 | UK | Bacteraemia | 2002 |
| EOE 3 | ST36 | UK | Bacteraemia | 2007 |
| EOE 23 | ST36 | UK | Bacteraemia | 2008 |
| EOE 30 | ST36 | UK | Bacteraemia | 1995 |
| EOE 35 | ST36 | UK | Bacteraemia | 1999 |
| EOE 41 | ST36 | UK | Bacteraemia | 2001 |
| EOE 42 | ST36 | UK | Bacteraemia | 2001 |
| EOE 45 | ST36 | UK | Bacteraemia | 2001 |
| EOE 52 | ST36 | UK | Bacteraemia | 1995 |
| EOE 54 | ST36 | UK | Bacteraemia | 1998 |
| EOE 57 | ST36 | UK | Bacteraemia | 1996 |
| EOE 61 | ST36 | UK | Bacteraemia | 1994 |
| EOE 65 | ST36 | UK | Bacteraemia | 1994 |
| EOE 72 | ST36 | UK | Bacteraemia | 1994 |
| EOE 73 | ST36 | UK | Bacteraemia | 1994 |
| EOE 78 | ST36 | UK | Bacteraemia | 1995 |
| EOE 83 | ST36 | UK | Bacteraemia | 1995 |
| EOE 84 | ST36 | UK | Bacteraemia | 1995 |
| EOE 86 | ST36 | UK | Bacteraemia | 1995 |
| EOE 88 | ST36 | UK | Bacteraemia | 1995 |
| EOE 89 | ST36 | UK | Bacteraemia | 1995 |
| EOE 90 | ST36 | UK | Bacteraemia | 1995 |
| EOE 91 | ST36 | UK | Bacteraemia | 1995 |
| EOE 94 | ST36 | UK | Bacteraemia | 1995 |
| EOE 96 | ST36 | UK | Bacteraemia | 1995 |
| EOE 97 | ST36 | UK | Bacteraemia | 1996 |
| EOE 98 | ST36 | UK | Bacteraemia | 1996 |
| EOE 99 | ST36 | UK | Bacteraemia | 1996 |
| EOE 100 | ST36 | UK | Bacteraemia | 1996 |
| EOE 101 | ST36 | UK | Bacteraemia | 1996 |
| EOE 102 | ST36 | UK | Bacteraemia | 1996 |
| EOE 103 | ST36 | UK | Bacteraemia | 1996 |
| EOE 104 | ST36 | UK | Bacteraemia | 1996 |
| EOE 105 | ST36 | UK | Bacteraemia | 1996 |
| EOE 106 | ST36 | UK | Bacteraemia | 1996 |
| EOE 122 | ST36 | UK | Bacteraemia | 1996 |
| EOE 125 | ST36 | UK | Bacteraemia | 1996 |
| EOE 129 | ST36 | UK | Bacteraemia | 1996 |
| EOE 130 | ST36 | UK | Bacteraemia | 1996 |
| EOE 137 | ST36 | UK | Bacteraemia | 1996 |
| EOE 140 | ST36 | UK | Bacteraemia | 1997 |
| EOE 154 | ST36 | UK | Bacteraemia | 1998 |
| EOE 155 | ST36 | UK | Bacteraemia | 1998 |

| | | | | |
|---|---|---|---|---|
| EOE 158 | ST36 | UK | Bacteraemia | 1998 |
| EOE 118 | ST36 | UK | Bacteraemia | 1996 |
| EOE 198 | ST36 | UK | Bacteraemia | 1999 |
| EOE 205 | ST36 | UK | Bacteraemia | 1999 |
| EOE 220 | ST36 | UK | Bacteraemia | 2000 |
| EOE 120 | ST36 | UK | Bacteraemia | 1996 |
| EOE 225 | ST36 | UK | Bacteraemia | 2000 |
| EOE 233 | ST36 | UK | Bacteraemia | 2001 |
| EOE 234 | ST36 | UK | Bacteraemia | 2001 |
| EOE 237 | ST36 | UK | Bacteraemia | 2001 |
| EOE 268 | ST36 | UK | Bacteraemia | 2003 |
| EOE 269 | ST36 | UK | Bacteraemia | 2003 |
| EOE 274 | ST36 | UK | Bacteraemia | 2003 |
| EOE 275 | ST36 | UK | Bacteraemia | 2003 |
| EOE 276 | ST36 | UK | Bacteraemia | 2003 |
| EOE 29 | ST36 | UK | Bacteraemia | 1995 |
| EOE 161 | ST36 | UK | Bacteraemia | 1998 |
| EOE 162 | ST36 | UK | Bacteraemia | 1998 |
| EOE 163 | ST36 | UK | Bacteraemia | 1998 |
| EOE 165 | ST36 | UK | Bacteraemia | 1998 |
| EOE 166 | ST36 | UK | Bacteraemia | 1998 |
| EOE 167 | ST36 | UK | Bacteraemia | 1998 |
| EOE 169 | ST36 | UK | Bacteraemia | 1998 |
| EOE 171 | ST36 | UK | Bacteraemia | 1998 |
| EOE 173 | ST36 | UK | Bacteraemia | 1998 |
| EOE 174 | ST36 | UK | Bacteraemia | 1998 |
| EOE 175 | ST36 | UK | Bacteraemia | 1998 |
| EOE 176 | ST36 | UK | Bacteraemia | 1998 |
| EOE 208 | ST36 | UK | Bacteraemia | 1998 |
| EOE 126 | ST36 | UK | Bacteraemia | 2006 |
| EOE 229 | ST36 | UK | Bacteraemia | 2001 |
| 2A8 | ST239 | Czech Republic | not recorded | 2001 |
| 2HK | ST239 | Czech Republic | not recorded | 2001 |
| 3HK | ST239 | Czech Republic | not recorded | 2000 |
| AGT1 | ST239 | Argentina | not recorded | 1997 |
| AGT120 | ST239 | Argentina | not recorded | 1998 |
| AGT67 | ST239 | Argentina | not recorded | 1997 |
| AGT9 | ST239 | Argentina | not recorded | 1997 |
| ANS46 | ST239 | Australia | not recorded | 1982 |
| BK2421 | ST239 | USA | not recorded | 1996 |
| BRA2 | ST239 | Brazil | not recorded | 1997 |
| BRA36 | ST239 | Brazil | not recorded | 1997 |
| CHI59 | ST239 | China | not recorded | 1998 |
| CHI61 | ST239 | China | not recorded | 1998 |
| CHL1 | ST239 | Chile | not recorded | 1997 |
| CHL151 | ST239 | Chile | not recorded | 1998 |

| | | | | |
|---|---|---|---|---|
| D71 | ST239 | Germany | Wound | 1996 |
| DEN907 | ST239 | Denmark | not recorded | 2001 |
| DEU17 | ST239 | Turkey | Blood | 2008 |
| DEU20 | ST239 | Turkey | Blood | 2008 |
| DEU29 | ST239 | Turkey | Blood | 2007 |
| DEU37 | ST239 | Turkey | Blood | 2007 |
| DEU9 | ST239 | Turkey | Aspiration | 2009 |
| ES26 | ST239 | Spain | Skin | 1996 |
| FRICAR | ST239 | France | not recorded | not recorded |
| GRE108 | ST239 | Greece | not recorded | 1998 |
| GRE4 | ST239 | Greece | not recorded | 1998 |
| H202 | ST239 | Thailand | Wound | 2006 |
| H211 | ST239 | Denmark | Lung | 2006 |
| H216 | ST239 | Denmark | Blood | 2006 |
| H482 | ST239 | Romania | Nose (carriage) | 1996 |
| HDG2 | ST239 | Portugal | not recorded | 1992 |
| HGSA142 | ST239 | Portugal | not recorded | 2003 |
| HGSA6 | ST239 | Portugal | not recorded | 1997 |
| HGSA9 | ST239 | Portugal | not recorded | 1997 |
| HSA10 | ST239 | Portugal | not recorded | 1992 |
| HSI216 | ST239 | Portugal | not recorded | 1997 |
| HU106 | ST239 | Hungary | not recorded | 1996 |
| HU109 | ST239 | Hungary | not recorded | 1996 |
| HU11 | ST239 | Turkey | Blood | 2007 |
| HU13 | ST239 | Turkey | Brain abscess | 2006 |
| HU16 | ST239 | Turkey | Spinal Fluid | 2007 |
| HU25 | ST239 | Brazil | not recorded | 1993 |
| HU5 | ST239 | Turkey | Catheter | 2006 |
| HU6 | ST239 | Turkey | Sputum | 2006 |
| HU7 | ST239 | Turkey | Abscess | 2007 |
| HU8 | ST239 | Turkey | Abscess | 2006 |
| ICP5011 | ST239 | Portugal | not recorded | 1993 |
| ICP5014 | ST239 | Portugal | not recorded | 1993 |
| ICP5062 | ST239 | Portugal | not recorded | 1993 |
| IU10 | ST239 | Turkey | Blood | 2007 |
| IU12 | ST239 | Turkey | Abscess | 2007 |
| IU17 | ST239 | Turkey | not recorded | 2007 |
| IU20 | ST239 | Turkey | not recorded | 2007 |
| IU4 | ST239 | Turkey | Sputum | 2006 |
| IU9 | ST239 | Turkey | Nasal swab | 2007 |
| LIT2 | ST239 | Lithuania | not recorded | not recorded |
| LIT68 | ST239 | Lithuania | Wound | 1996 |
| LIT76 | ST239 | Lithuania | Wound | 1996 |
| LIT89 | ST239 | Lithuania | Blood | 1996 |
| M116 | ST239 | Vitnam | not recorded | 2004 |
| M1229 | ST239 | Denmark | Lung | 2009 |

| | | | | |
|---|---|---|---|---|
| M278 | ST239 | Portugal | Nose (carriage) | 2005 |
| M418 | ST239 | India | Nose (carriage) | 2006 |
| M705 | ST239 | Thailand | Lung | 2007 |
| M74 | ST239 | Extensive travel | not recorded | not recorded |
| M996 | ST239 | China | Lung | 2008 |
| MAL1 | ST239 | Malaysia | Wound | 1996 |
| MAL11 | ST239 | Malaysia | Wound | 1996 |
| MAL119 | ST239 | Malaysia | not recorded | 1996 |
| MAL215 | ST239 | Malaysia | not recorded | not recorded |
| MAL3 | ST239 | Malaysia | not recorded | not recorded |
| MAL35 | ST239 | Malaysia | not recorded | not recorded |
| MU11 | ST239 | Turkey | Blood | 2006 |
| Na21 | ST239 | Sri Lanka | not recorded | 1996 |
| P32 | ST239 | Poland | Lung | 1996 |
| R3J | ST239 | Poland | not recorded | not recorded |
| RA6 | ST239 | Argentina | Skin | 1996 |
| RA7 | ST239 | Argentina | Wound | 1996 |
| TUR1 | ST239 | Turkey | not recorded | 1996 |
| TUR27 | ST239 | Turkey | not recorded | 1996 |
| TUR9 | ST239 | Turkey | not recorded | 1995 |
| TW20 | ST239 | UK | not recorded | 2003 |
| UCO159 | ST239 | Argentina | not recorded | not recorded |
| UK102 | ST239 | UK | not recorded | 1996 |
| UK105 | ST239 | UK | not recorded | not recorded |
| URU110 | ST239 | Uruguay | not recorded | 1998 |
| URU34 | ST239 | Uruguay | not recorded | 1997 |

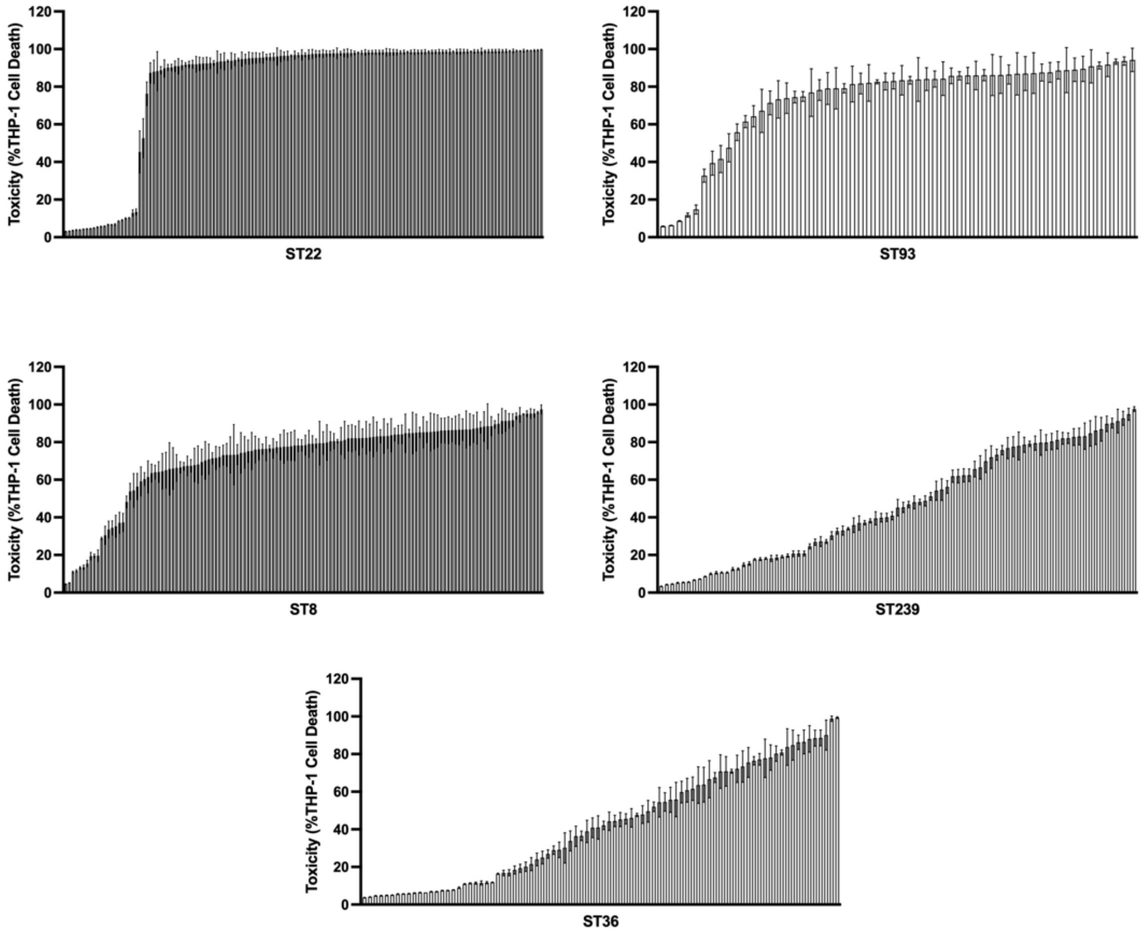


**Supplementary Figure 1: Toxicity of 475 MRSA clinical isolates**

The toxicity of each clinical isolate from five MRSA multi-locus sequence types (STs) (ST22 (n=110), ST93 (n=58), ST8 (n=134), ST239 (n=87) and ST36 (n=86)) was examined by incubating culture supernatant (either diluted to 30% using sterile TS broth (ST22, ST93 and ST8) or used neat (100%; ST239 and ST36)) with cultured THP-1 cells and toxicity as a measure of THP-1 cell death determined using flow cytometry. Each bar represents the mean of one clinical isolate quantified using three biological repeats with error bars representing the standard deviation.

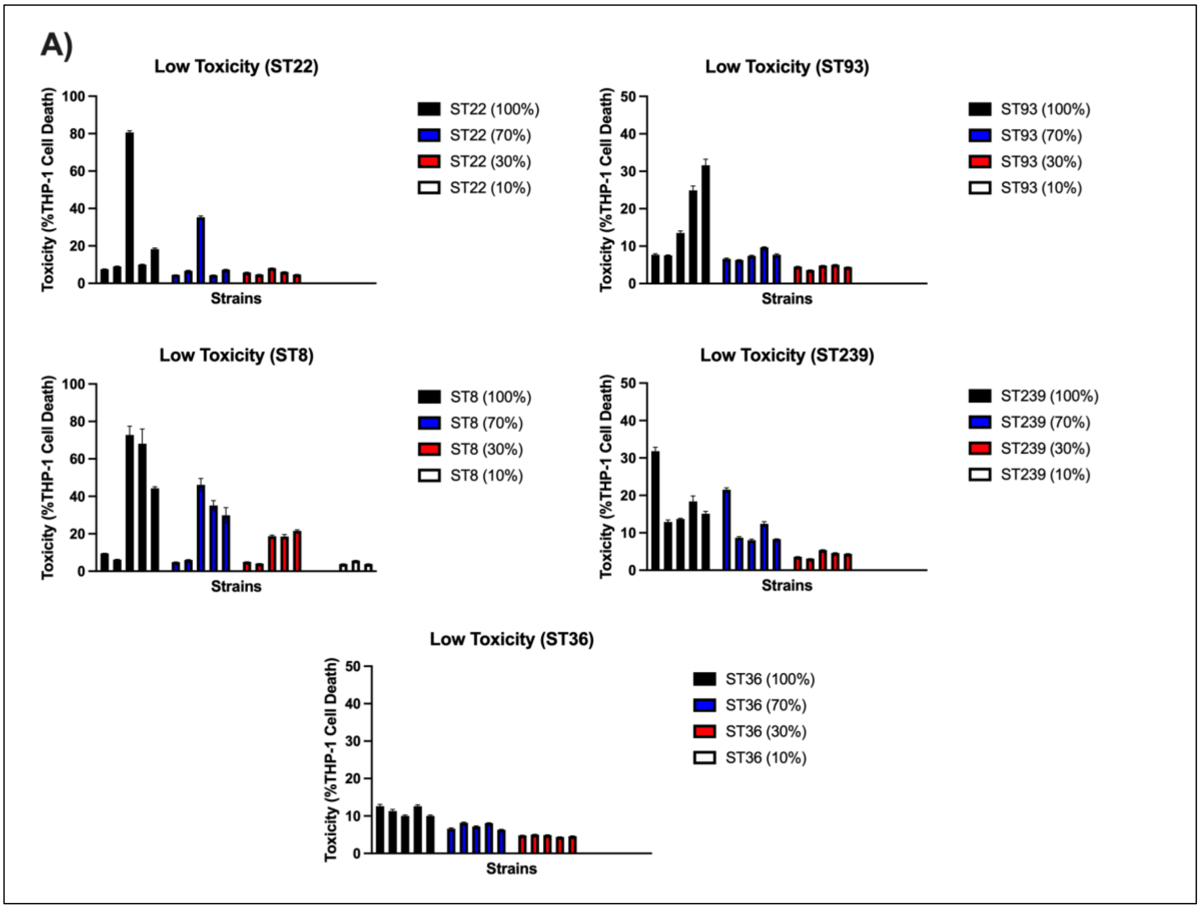
A)
Low Toxicity (ST22)
Toxicity (%THP-1 Cell Death)
Strains
ST22 (100%)
ST22 (70%)
ST22 (30%)
ST22 (10%)
Low Toxicity (ST93)
ST93 (100%)
ST93 (70%)
ST93 (30%)
ST93 (10%)
Low Toxicity (ST8)
ST8 (100%)
ST8 (70%)
ST8 (30%)
ST8 (10%)
Low Toxicity (ST239)
ST239 (100%)
ST239 (70%)
ST239 (30%)
ST239 (10%)
Low Toxicity (ST36)
ST36 (100%)
ST36 (70%)
ST36 (30%)
ST36 (10%)

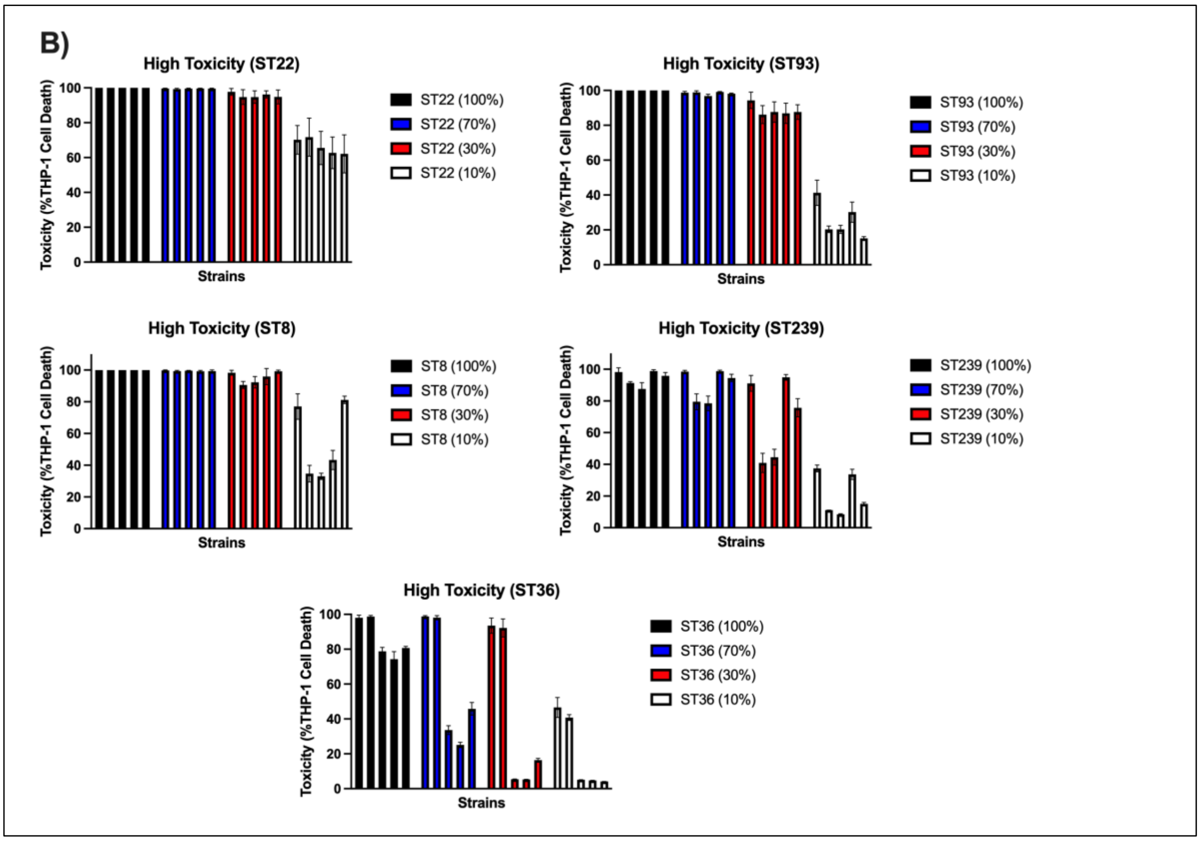
B)
High Toxicity (ST22)
Toxicity (%THP-1 Cell Death)
Strains
ST22 (100%)
ST22 (70%)
ST22 (30%)
ST22 (10%)
High Toxicity (ST93)
ST93 (100%)
ST93 (70%)
ST93 (30%)
ST93 (10%)
High Toxicity (ST8)
ST8 (100%)
ST8 (70%)
ST8 (30%)
ST8 (10%)
High Toxicity (ST239)
ST239 (100%)
ST239 (70%)
ST239 (30%)
ST239 (10%)
High Toxicity (ST36)
ST36 (100%)
ST36 (70%)
ST36 (30%)
ST36 (10%)

**Supplementary Figure 2: Supernatant dilutions of the lowest and highest toxic isolates from each of the five MRSA sequence types.**

Cell-free culture supernatant of the five lowest **(A)** and highest **(B)** toxic isolates from each of the five MRSA STs were diluted to either 10%, 30% or 70% supernatant using sterile TS broth or used neat (100%), and toxicity determined as a measure of THP-1 cell death. The toxicity of each isolate was quantified using three biological repeats with each bar representing the mean of individual isolates and error bars indicating the standard deviation.
